# The relative transmissibility of shigellosis among male and female individuals in Hubei Province, China: a modelling study

**DOI:** 10.1101/677088

**Authors:** Zeyu Zhao, Qi Chen, Bin Zhao, Mikah Ngwanguong Hannah, Ning Wang, Yuxin Wang, Xianfa Xuan, Jia Rui, Meijie Chu, Yao Wang, Xingchun Liu, An Ran, Lili Pan, Yi-Chen Chiang, Yanhua Su, Benhua Zhao, Tianmu Chen

## Abstract

**Objective:** Shigellosis has been a heavy burden in China. However, its relative transmissibility in male and female individuals remains unclear.

**Method:** A sex-based Susceptible–Exposed–Infectious/Asymptomatic–Recovered (SEIAR) model was applied to explore the dataset of reported shigellosis cases built by Hubei Province from 2005 to 2017. Two indicators, secondary attack rate (SAR) and relative ratio of SAR between males and females, were developed to assess the relative transmissibility in males and females.

**Results:** The number of cases and reported incidences in males and females demonstrated a significant decreasing trend (Male trend: *χ*^2^ = 11.268, *P* = 0.001, Female trend: *χ*^2^ = 11.144, *P* = 0.001). SEIAR model had a great fitting effect with the data of shigellosis (*P* < 0.001). Our simulation revealed that, when parameter *β_fm_* = 0, the greatest decrease in cases were obtained for different genders. The median value for *SAR_mm_* was 2.3225 × 10^−8^ (Range: 1.7574 × 10^−8^ – 3.8565 × 10^−8^), *SAR_mf_* was 2.5729 × 10^−8^ (Range: 1.3772 × 10^−8^ – 3.2773 × 10^−8^), *SAR_fm_* was 2.7630 × 10^−8^ (Range: 1.8387 × 10^−8^ – 4.2638 × 10^−8^) and *SAR_ff_* was 2.1061 × 10^−8^ (Range: 1.0201 × 10^−8^ – 3.2140 × 10^−8^). The median value of relative ratio calculated by *SAR* in *mm* versus (vs) *mf* was 0.93 (Range: 0.75 – 1.47), *mm* vs *fm* was 0.90 (Range: 0.41 – 1.81), *mm* vs *ff* was 1.07 (Range: 0.55 – 2.93), *mf* vs *fm* was 0.99 (Range: 0.32 – 1.25), *mf* vs *ff* was 1.17 (Range: 0.43 – 3.21) and *ff* vs *fm* was 0.75 (Range: 0.35 – 1.06).

**Conclusion:** Transmissibility of shigellosis is different among male and female individuals. Shigellosis seems to be more transmissible in males than in females.

**Author summary:** Shigellosis, also known as bacillary dysentery, is an infectious disease caused by the genus *Shigella spp*. Developing countries have high disease burden of shigellosis. However, its relative transmissibility in male and female individuals remains unclear. In this study, we employed a mathematical model to explore the dataset of reported shigellosis cases built by Hubei Province, China from 2005 to 2017. Two indicators, secondary attack rate (SAR) and relative ratio of SAR between males and females, were developed to assess the relative transmissibility in males and females. We found that shigellosis has medium transmissibility among male and female individuals. The disease seems to be more transmissible in males than in females.

## Introduction

Shigellosis, also known as bacillary dysentery, is an infectious disease caused by the genus *Shigella spp*, and often occurs in summer and autumn. *Shigella flexneri* is the main cause of endemic diarrhoea in low and middle income countries, and lays a heavy burden on these countries, especially in children aged 1-4 years old [1]. According to the Chinese Center for Disease Control and Prevention, about 150,000~450,000 cases were reported annually within the period 2005 to 2014 [2]. Although there has been an improvement in the quality of water and sanitaion, shigellosis remains a major public health problem in some developing countries, including China [3, 4].

Bacillary dysentery is an intestinal infectious disease, which can be transmitted via the consumption of contaminated food or water [5]. Humans are the only natural host for *shigella*. Shigellosis has low infectious dose and transmission primarily occurs from person-to-person [1]. Previously, a study was done [6] in China during which it was evaluated that, contaminated water and food hardly contributed to shigellosis. According to these reports, the incidence rate of bacillary dysentery is higher in males than in females [7]. So, was there a shift route in the transmission of shigellosis in developing countries? What is the process of transmission among individuals? What caused the different incidence in males and females?

The distribution of time and space were focused more in the model studies of shigellosis, while population-based research was less [8–12]. Studies have showed that the Susceptible–Exposed–Infectious/Asymptomatic–Recovered–Water (SEIARW) model has a great fitting effect [6]. However, it does not estimate the transmissibility of bacillary dysentery between males and females. Therefore, a new model–SEIAR, is formed by simplifying the SEIARW model. We adopted secondary attack rate (*SAR*) to quantify the infectivity of shigellosis and relative ratio to assess the transmissibility of shigellosis between males and females.

In this study, we collected Shigellosis cases reported in Hubei Province, China, adopted SEIAR model to fit the data, calculated related index and figured out the transmissibility of shigellosis between males and females.

## Materials and Methods

### Data sources

A dataset of shigellosis reported cases built by Hubei Province from January 2005 to December 2017, were collected from China Information System for Disease Control and Prevention. We cleared up the date and sex (male or female) of onset of illness for each case. The informations of the population such as, birth rate, death rate and total population were obtained from Hubei Statistical Yearbook.

### Shigellosis model between different genders

According to a new review [1], the transmission of shigellosis is mainly from person-to-person in developed countries. Therefore, the SEIAR model was developed according to the natural history of shigellosis between male and female individuals (Figure 1). The pattern followed by the model was from person-to-person, which contained susceptible (S), exposed (E), symptomatic (I), asymptomatic (A), and recovered (R) individuals (Table 1). We used the subscript *m* to represent male and *f* to represent female. In the model, we assumed that:

a. Relative rate of transmission among male and female individuals was *β_m_* and *β_f_*, respectively;
b. Relative rate of transmission from male to female was *β_mf_* and from female to male was *β_fm_*.

**Figure 1.**
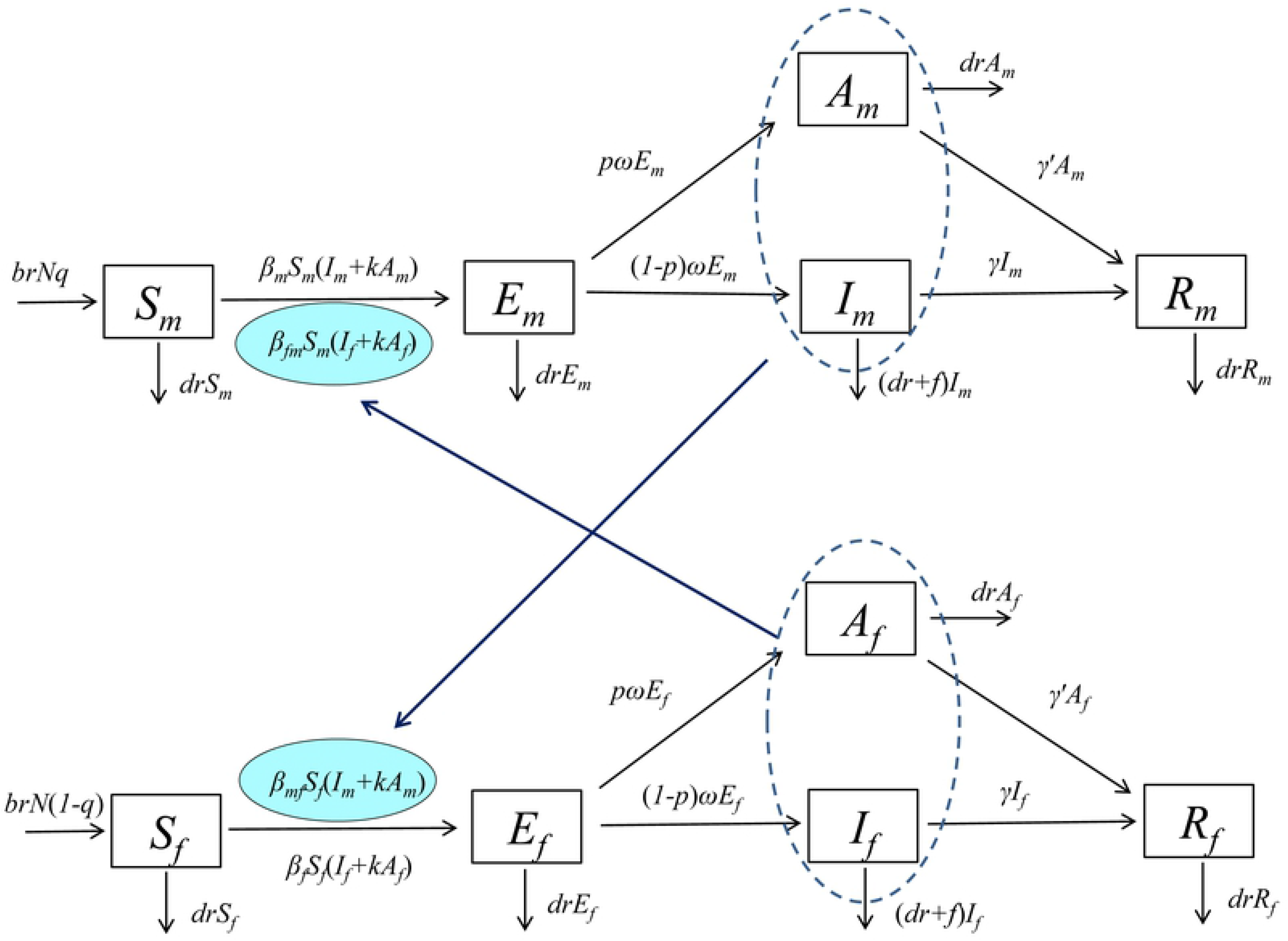
Flowchart of transmission SEIAR model of shigellosis in different genders.

**Table 1.**
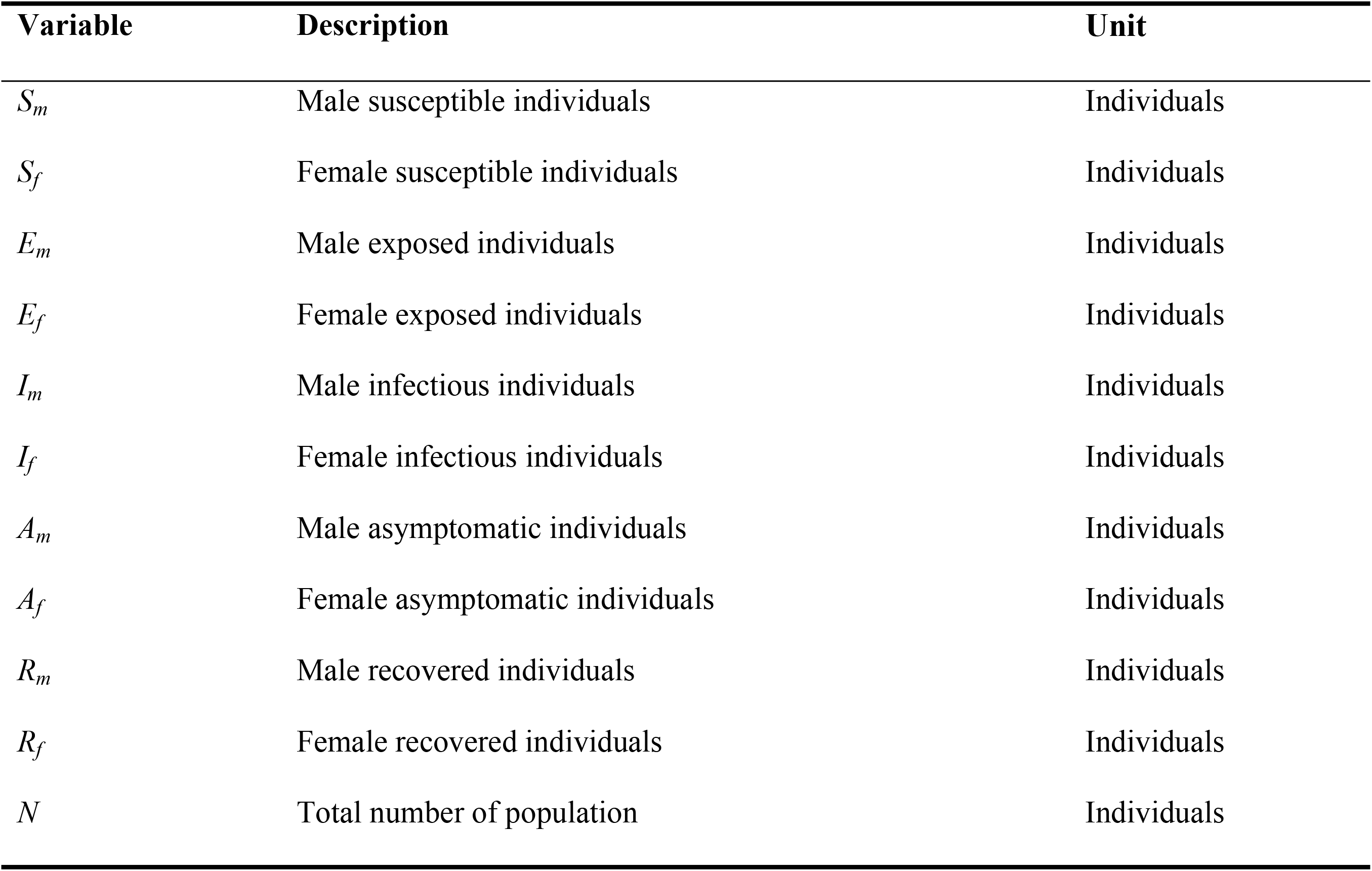
Variables with the intersex transmission SEIAR model

**Table 2.**
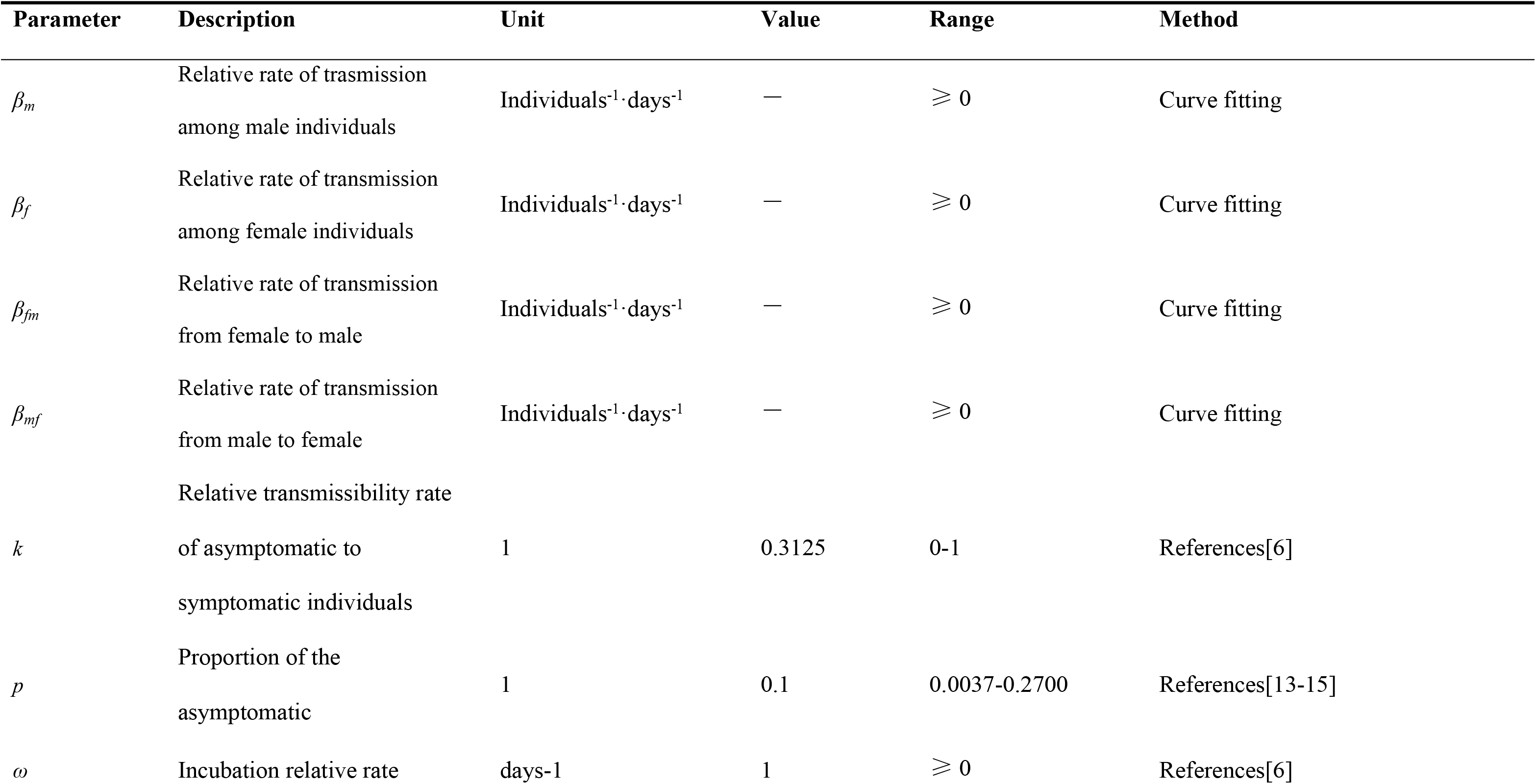
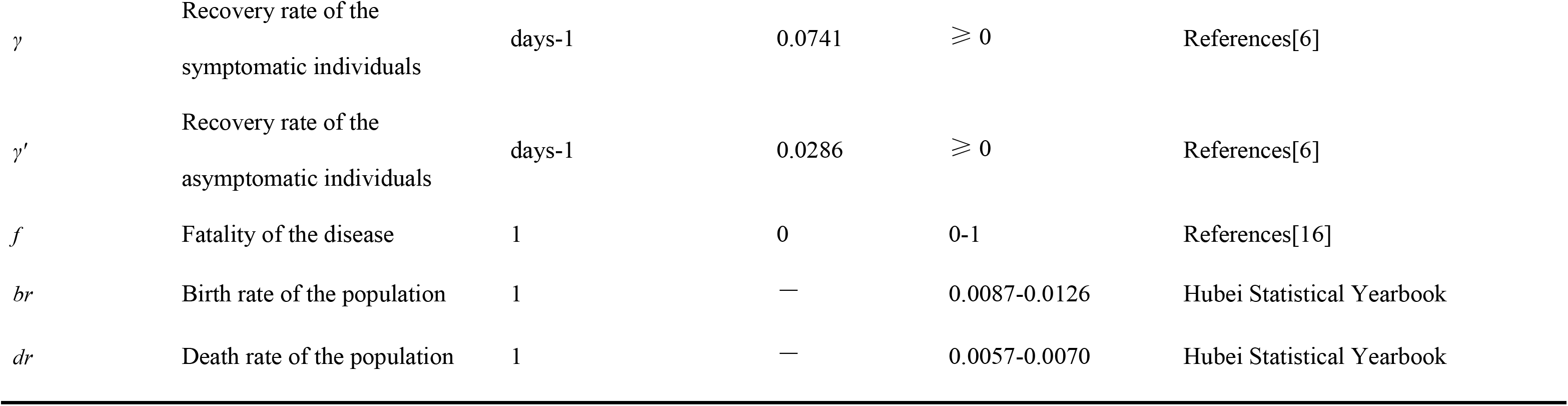
Parameter description and values of SEIAR model

| Parameter | Description | Unit | Value | Range | Method |
| --- | --- | --- | --- | --- | --- |
| $\beta_m$ | Relative rate of trasmission<br>among male individuals | Individuals<sup>-1</sup>·days<sup>-1</sup> | — | $\geq 0$ | Curve fitting |
| $\beta_f$ | Relative rate of transmission<br>among female individuals | Individuals<sup>-1</sup>·days<sup>-1</sup> | — | $\geq 0$ | Curve fitting |
| $\beta_{fm}$ | Relative rate of transmission<br>from female to male | Individuals<sup>-1</sup>·days<sup>-1</sup> | — | $\geq 0$ | Curve fitting |
| $\beta_{mf}$ | Relative rate of transmission<br>from male to female | Individuals<sup>-1</sup>·days<sup>-1</sup> | — | $\geq 0$ | Curve fitting |
| $k$ | Relative transmissibility rate<br>of asymptomatic to<br>symptomatic individuals | 1 | 0.3125 | 0-1 | References[6] |
| $p$ | Proportion of the<br>asymptomatic | 1 | 0.1 | 0.0037-0.2700 | References[13-15] |
| $\omega$ | Incubation relative rate | days-1 | 1 | $\geq 0$ | References[6] |
| $\gamma$ | Recovery rate of the symptomatic individuals | days-1 | 0.0741 | $\geq 0$ | References[6] |
| $\gamma'$ | Recovery rate of the asymptomatic individuals | days-1 | 0.0286 | $\geq 0$ | References[6] |
| $f$ | Fatality of the disease | 1 | 0 | 0-1 | References[16] |
| $br$ | Birth rate of the population | 1 | — | 0.0087-0.0126 | Hubei Statistical Yearbook |
| $dr$ | Death rate of the population | 1 | — | 0.0057-0.0070 | Hubei Statistical Yearbook |

**Table 3.** *R^2^* of model and reported cases in different genders from 2005 to 2017 in Hubei Province, China

| Year | Male |  | Female |  |
| --- | --- | --- | --- | --- |
| | $R^2$ | $p$ | $R^2$ | $p$ |
| 2005 | 0.989 | < 0.001 | 0.991 | < 0.001 |
| 2006 | 0.995 | < 0.001 | 0.992 | < 0.001 |
| 2007 | 0.992 | < 0.001 | 0.987 | < 0.001 |
| 2008 | 0.984 | < 0.001 | 0.986 | < 0.001 |
| 2009 | 0.982 | < 0.001 | 0.984 | < 0.001 |
| 2010 | 0.989 | < 0.001 | 0.982 | < 0.001 |
| 2011 | 0.985 | < 0.001 | 0.982 | < 0.001 |
| 2012 | 0.989 | < 0.001 | 0.979 | < 0.001 |
| 2013 | 0.977 | < 0.001 | 0.983 | < 0.001 |
| 2014 | 0.986 | < 0.001 | 0.983 | < 0.001 |
| 2015 | 0.977 | < 0.001 | 0.965 | < 0.001 |
| 2016 | 0.985 | < 0.001 | 0.988 | < 0.001 |
| 2017 | 0.986 | < 0.001 | 0.978 | < 0.001 |

We assumed that in both males and females:

a. The disease does not spread vertically, and the individuals born in various groups are all susceptible. The natural birth rate is *br*, and the natural mortality rate is *dr*;
b. The proportion of latency patients (*1-p*)*E* (0 ≤ *p* ≤ 1) will change to infected person *I* after one incubation period, while another part of the latent *pE* will become a latent infected person *A* after an incubation period. Therefore, at time *t*, the speed from *E* to *I* is proportional to the latency group, the proportional coefficient is (*1-p*)*ω*, and the speed from *E* to *A* is proportional to the latency population, and the proportional coefficient is *pω*;
c. The speeds removed from *I* and *A* are proportional to the number of people in both groups, and the proportional coefficients are *γ* and *γ*′, respectively.

The model was expressed as follows:

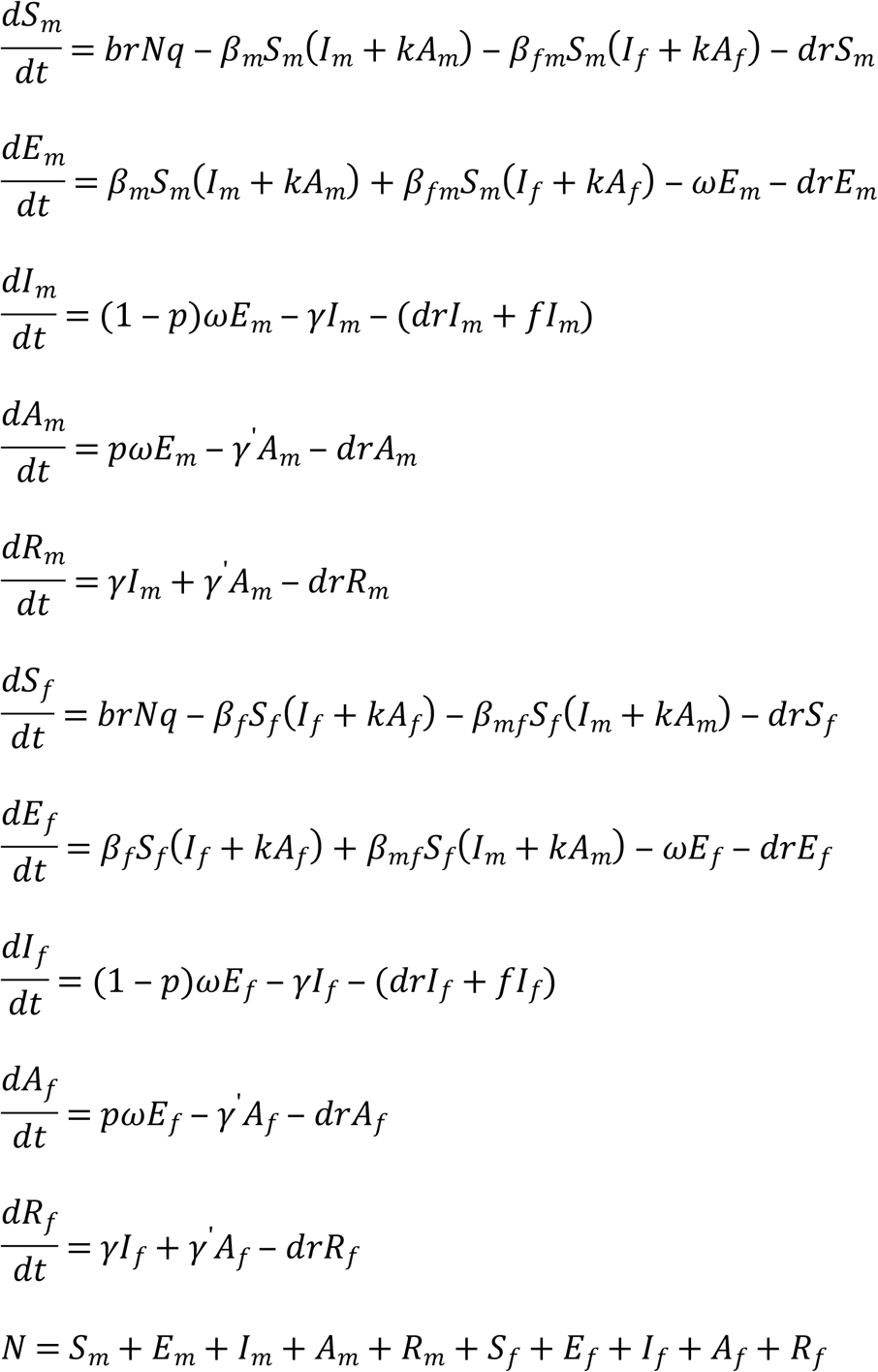

The left side of the equation shows the instantaneous rate of change of *S*, *E*, *I*, *A* and *R* at time *t*. In the equations, the parameters *β_m_*, *β_f_*, *β_mf_*, *β_fm_*, *k*, *ω*, *p*, *f*, *γ* and *γ*′ refer to relative rate of transmission in males, relative rate of transmission in females, relative rate of transmission from male to female, relative rate of transmission from female to male, relative transmissibility of asymptomatic to symptomatic individuals, incubation period, proportion of asymptomatic individuals, fatality of shigellosis, recovery rate of symptomatic individuals, recovery rate of asymptomatic individuals.

In the model, we quantify the transmissibility of shigellosis by secondary attack rate (*SAR*), which is defined as the probability that infection occurs among susceptible persons within a reasonable incubation period, following contact with an infectious person or an infectious source. Relative ratio of transmission is developed to assess the relative transmissibility of male versus female. We calculated the *SAR* and relative ratio by the equation as follows:

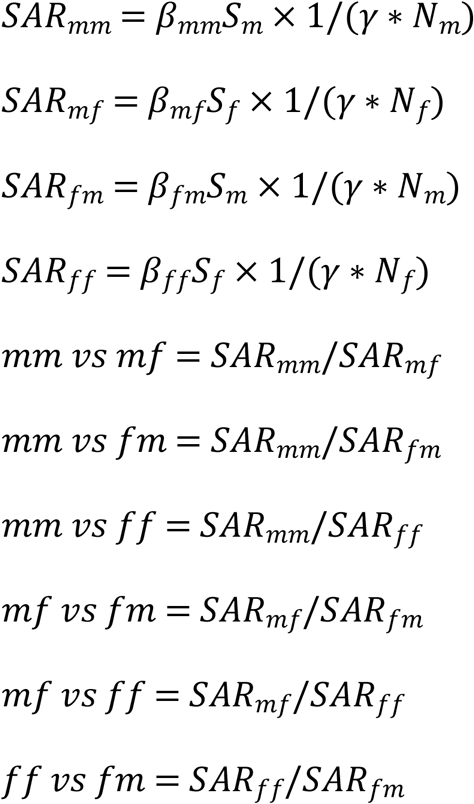

In the equation, we defined the subscripts; *mm*, *ff*, *mf* and *fm* as among males, among females, from female to male, and from male to female, respectively. *mm* vs *mf*, *mm* vs *fm*, *mm* vs *ff*, *mf* vs *fm*, *mf* vs *ff* and *ff* vs *fm* refer to *mm*, *ff*, *mf* and *fm*, which have similar definitions as with the subscript of *SAR*.

### Estimation of Parameters

According to epidemiological characteristics of shigellosis and previous study [6], we set *k*, *ω*, *γ* and *γ*′ as 0.3125, 1.0000, 0.0741 and 0.0286, respectively. The proportions of asymptomatic individuals were reported to range from 0.0037 to 0.2700 [13–15]. We set *p* = 0.1 in SEIAR model. The fatality rate of the disease reported in a study decreased from 0.00031 to 0.00088 from 1991 to 2000 [16]. Considering the fatality rate of shigellosis is extremely low, we set *f* = 0. The values of *β_m_*, *β_f_*, *β_mf_* and *β_fm_* were generated by curve fitting using SEIAR model and reported shigellosis data. In order to simulate the contribution of *β_m_*, *β_f_*, *β_mf_* and *β_fm_* during the transmission, we performed a “knock-out” simulation in five scenarios: A) *β_m_* = 0; B) *β_mf_* = 0; C) *β_f_* = 0; D) *β_fm_* = 0; E) control (no intervention).

### Simulation method and statistical analysis

Berkeley Madonna 8.3.18 (developed by Robert Macey and George Oster of the University of California at Berkeley. Copyright ©1993-2001 Robert I. Macey & George F. Oster) was employed for model simulation. Simulation methods were as previously described [6, 17–20]. Microsoft Office Excel 2007 (Microsoft, Redmond, WA, USA) and GraphPad Prism 7.00 (GraphPad Software, La Jolla California, USA) were employed for figure development and data analysis. SPSS 21.0 (IBM Corp, Armonk, NY, USA) was used to calculate coefficient of determination (*R*^2^) by curve fitting, which was adopted to judge the goodness of fit of the model.

### Ethics

This effort of disease control was part of CDC’s routine responsibility in Hubei Province; therefore, institutional review and informed consent were not required for this study.

## Results

### Epidemiological characteristics of shigellosis reported cases

From 2005 to 2017, 130770 shigellosis cases (including 73981male cases and 56789 female cases) were reported in Hubei province (Figure 2). The median of incidences reported annually was 21.68 per 100000 persons (range: 6.10 – 32.63 per 100000 persons) in males and 17.91 per 100000 persons (range: 5.87 – 26.51 per 100000 persons) in females. It demonstrated that, the number of cases and reported incidences in males and females had significantly decreased. (Male trend: *χ*^2^ = 11.268, *P* = 0.001, Female trend: *χ*^2^ = 11.144, *P* = 0.001).

**Figure 2.**
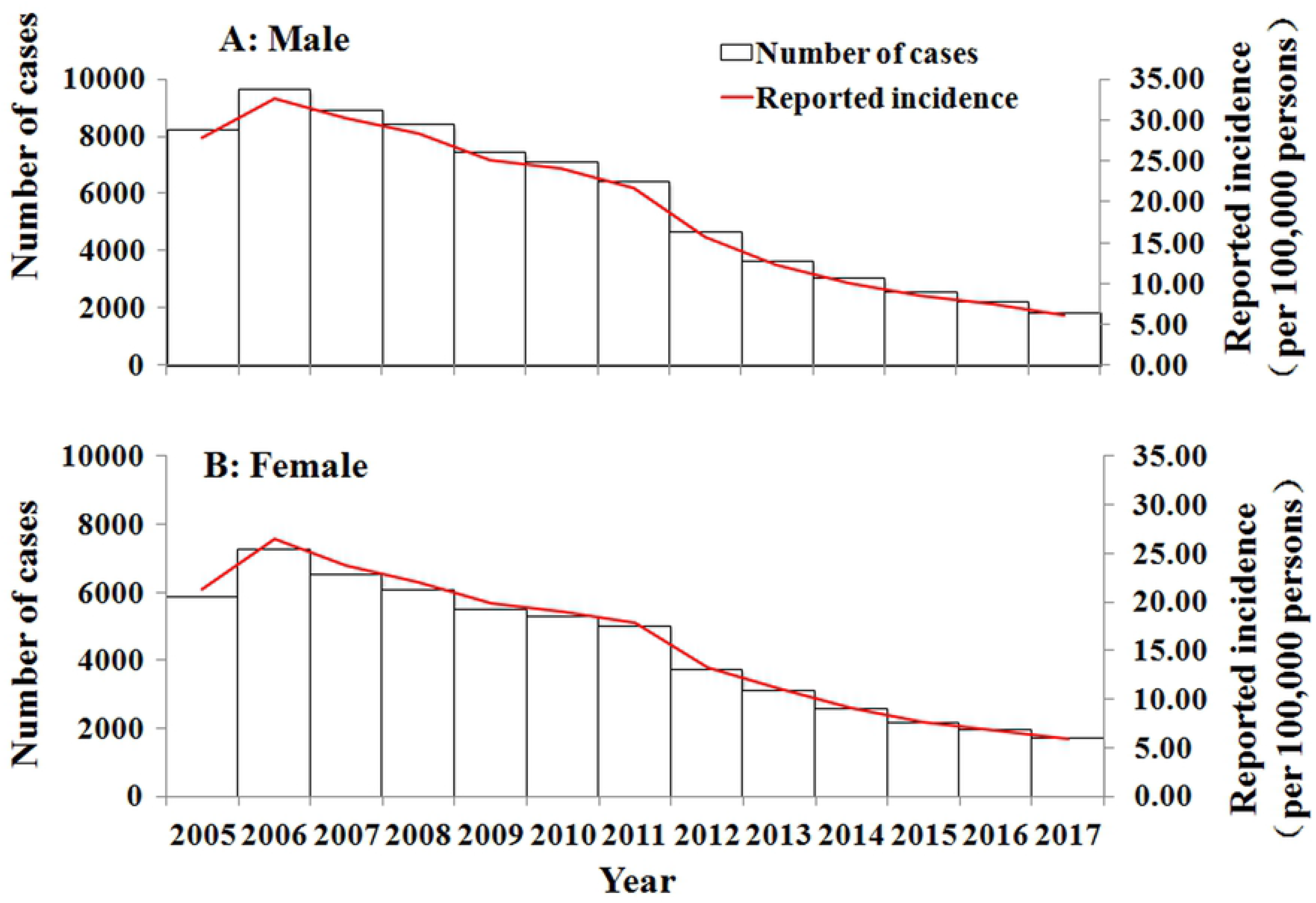
Reported cases and incidence of Shigellosis in different genders from 2005 to 2017 in Hubei province (A: Male; B: Female).

### Curve fitting results

The results of curve fitting showed that the SEIAR model fitted the data well (Figure 3). The *R*^2^ of SEIAR model of different genders each year were shown in Table 1. The model had a great fitting effect with the data of shigellosis (Supplementary Table 1).

**Figure 3.**
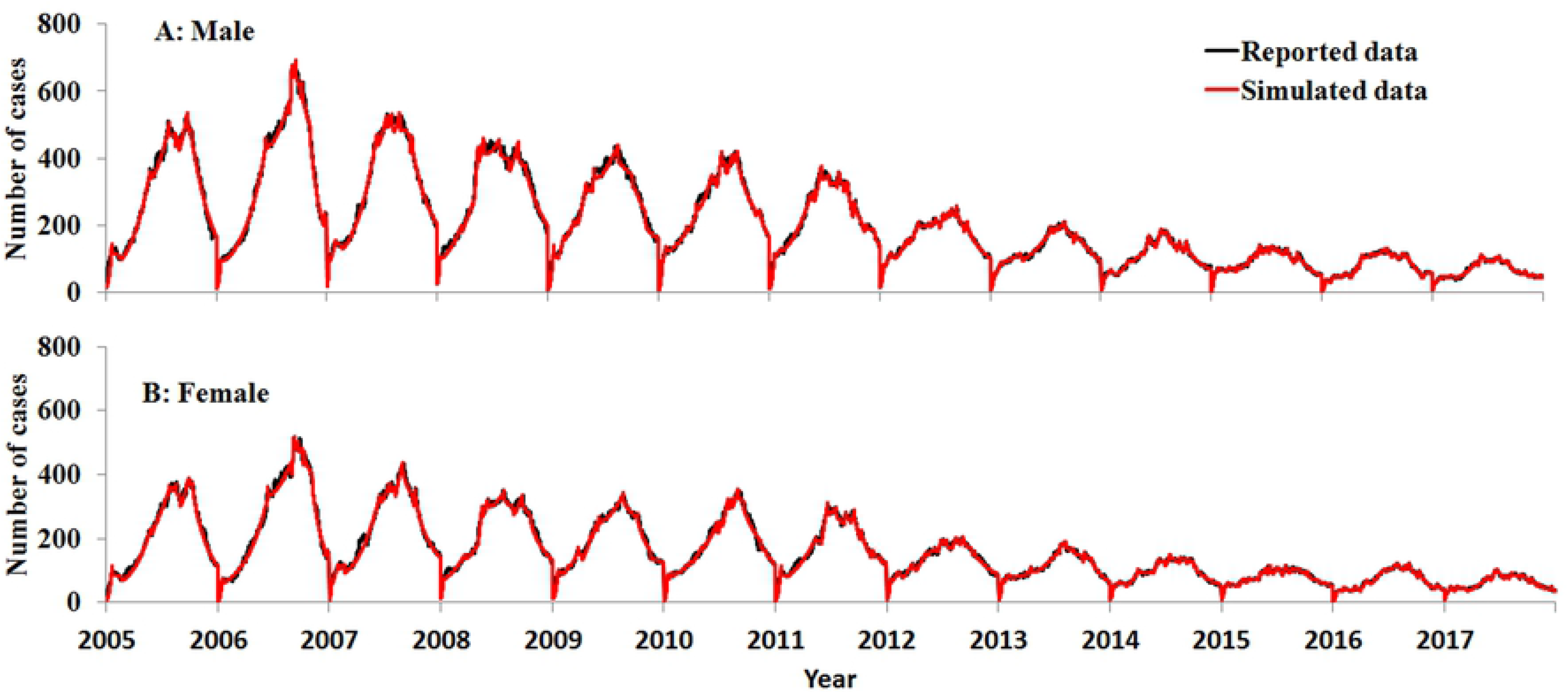
Curve fitting of Model to reported data in different genders from 2005 to 2017 in Hubei (A: Male; B: Female).

### The transmissibility of shigellosis

From Figure 4, the results of the “knock-out” simulation showed that the number of cases in different genders using parameters *β_m_* = 0, *β_f_* = 0, *β_mf_* = 0 and *β_fm_* = 0 were lower than in the control group. When *β_fm_* = 0, the number of cases decreased most in different genders.

**Figure 4.**
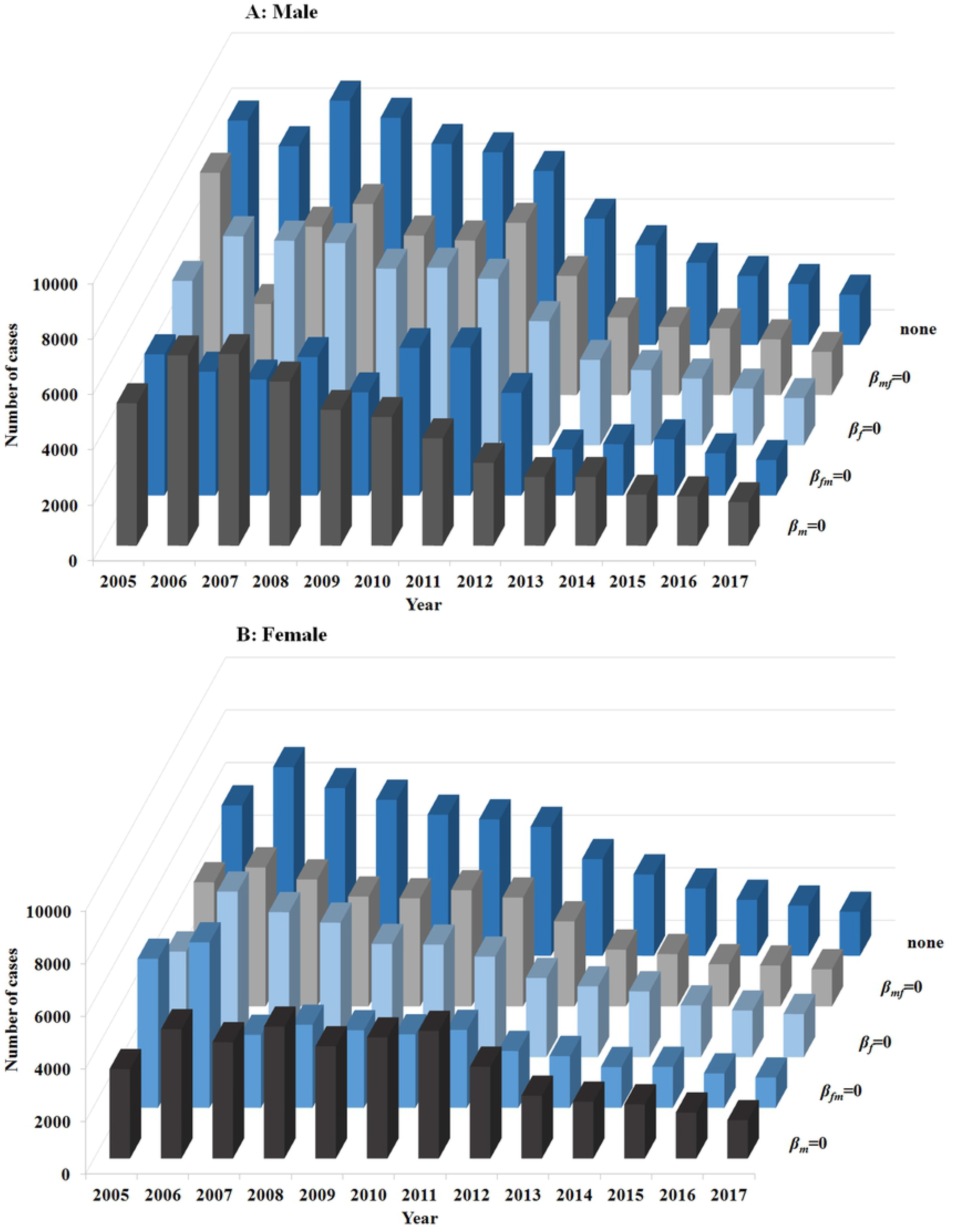
The results to simulate the contribution of *β* during the transmission in different genders (A: Male; B: Female; *β_m_* = 0, control transmission among male; *β_fm_* = 0, control transmission among female; *β_f_* = 0, control transmission from female to male; *β_mf_* = 0, control transmission from male to female; and control defined as “None”).

Figure 5 showed the difference between the mean and 95% confidence interval (*CI*) from 2005 to 2017 using *β_m_*, *β_f_*, *β_mf_* and *β_fm_*. The mean value when using *β_m_* was 1.9240 × 10^−9^ (95% *CI:* 1.6621 × 10^−9^ – 6.6121 × 10^−9^), using *β_f_* was 1.5645 × 10^−9^ (95% *CI:* 1.3521 × 10^−9^ – 1.7769 × 10^−9^), using *β_fm_* was 2.1572 × 10^−9^ (95% *CI:* 1.9159 × 10^−9^ – 2.3986 × 10^−9^) and using *β_mf_* was 1.8750 × 10^−9^ (95% *CI:* 1.6846 × 10^−9^ – 2.0654 × 10^−9^).

**Figure 5.**
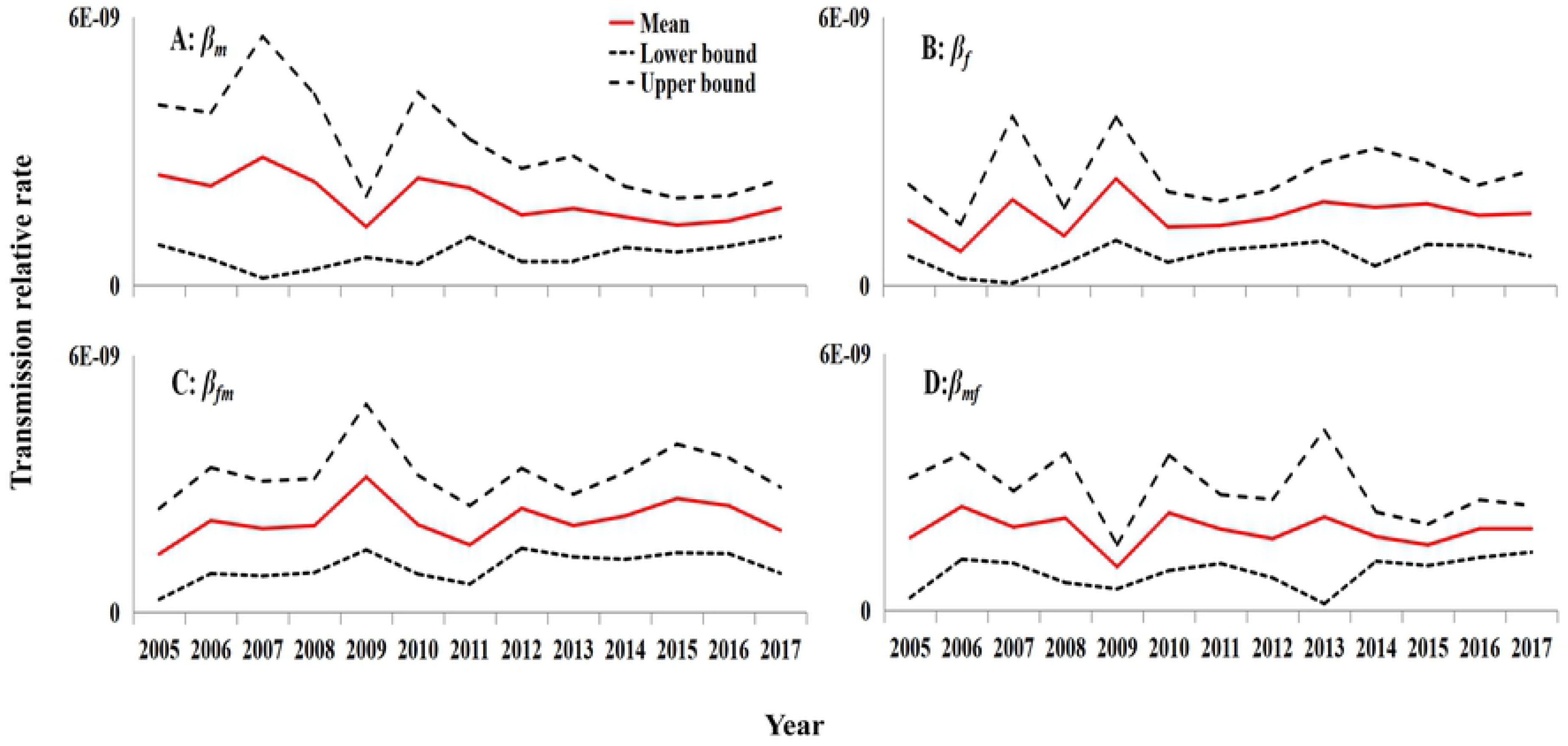
The parameter of *β_m_, *β_f_*, β_mf_* and *β_fm_* during the transmission from 2005 to 2017 in Hubei (A: *β_m_*, transmission relative rate among male; B: *β_f_*, transmission relative rate among female; C: *β_mf_*, transmission relative rate from male to female; D: *β_fm_*, transmission relative rate from male to female).

The results of *SAR* from 2005 to 2017 were showed in Figure 6 and Figure 7. The median value of *SAR_fm_* was 2.3225 × 10^−8^ (Range: 1.7574 × 10^−8^ – 3.8565 × 10^−8^). The median value of *SAR_mf_* was 2.5729 × 10^−8^ (Range: 1.3772 × 10^−8^ – 3.2773 × 10^−8^). The median value of *SAR_fm_* was 2.7630 × 10^−8^ (Range: 1.8387 × 10^−8^ – 4.2638 × 10^−8^). The median value of *SAR_ff_* was 2.1061 × 10^−8^ (Range: 1.0201 × 10^−8^ – 3.2140 × 10^−8^).

**Figure 6.**
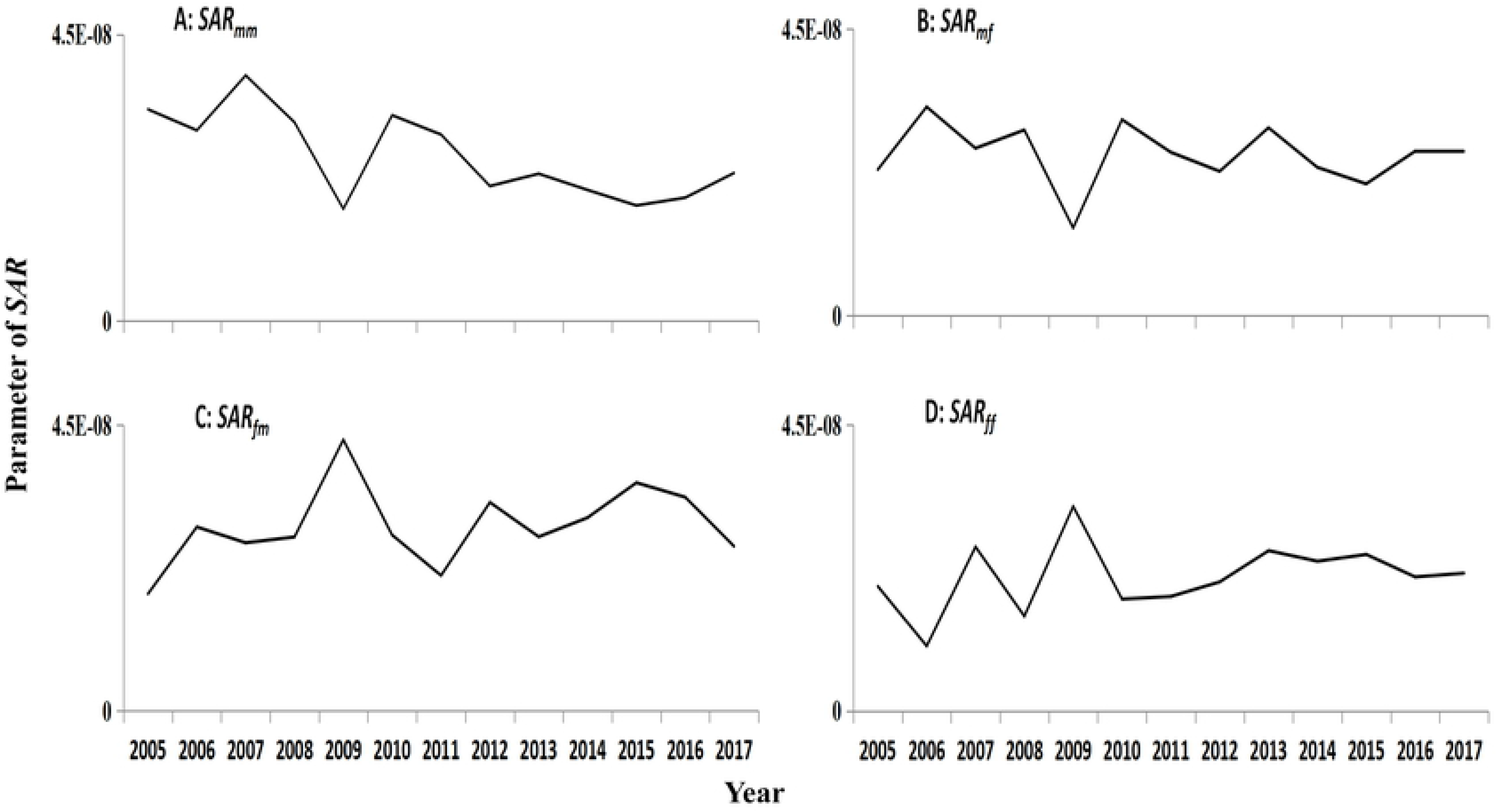
The *SAR_mm_*, *SAR_mf_*, *SAR_fm_* and *SAR_ff_* estimated by Model from 2005 to 2017 in Hubei (A: *mm*, among male; B: *mf*, from male to female; C: *fm*, from female to male; D: *ff*, among female).

**Figure 7.**
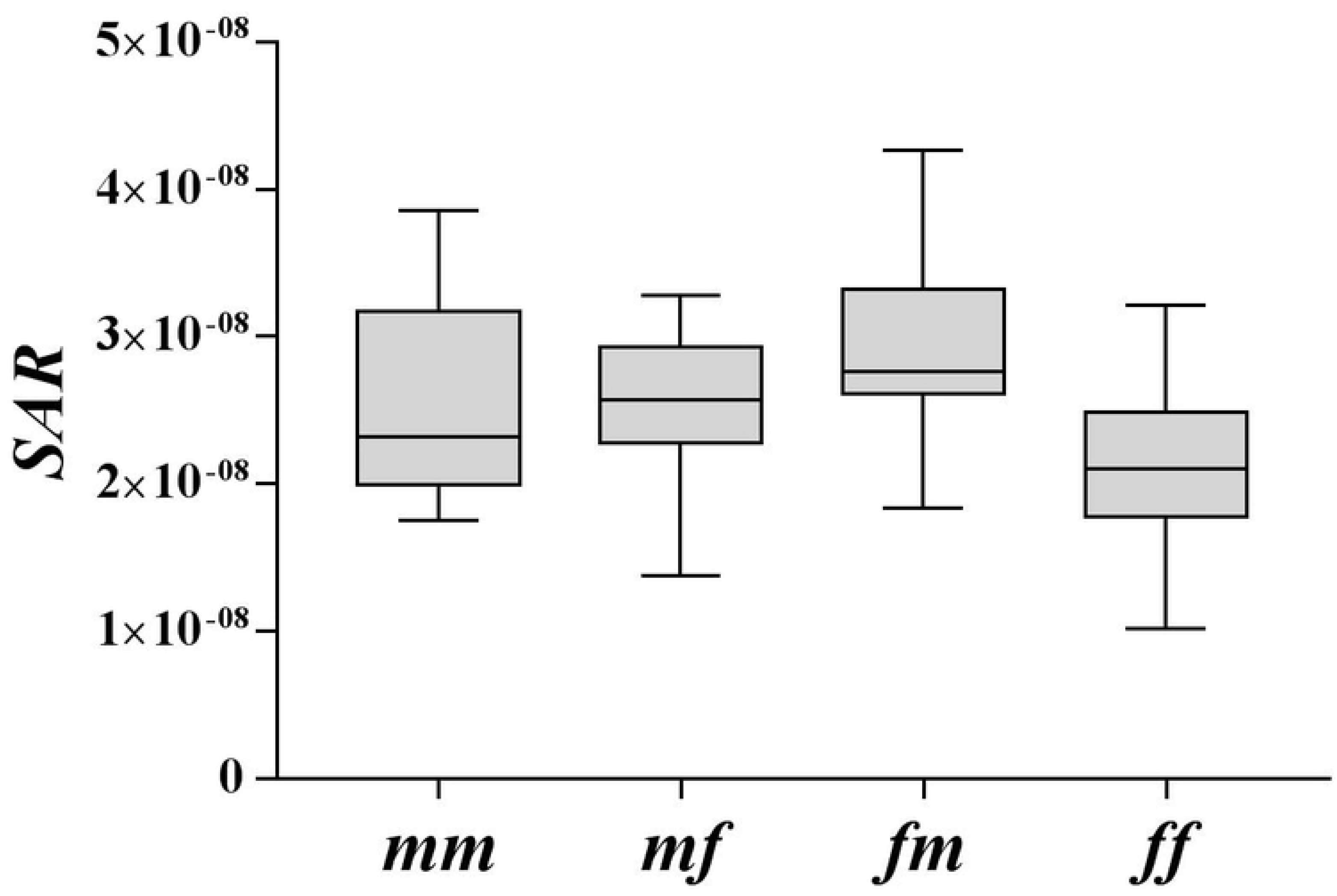
Box-plot of SAR from 2005 to 2017 in Hubei (*mm*: among male; *ff:* among female; *fm:* from female to male; *mf*: from male to female).

The results of relative ratio of the dataset were depicted in Figure 8. The median value of relative ratio calculated by *SAR* in *mm* vs *mf* was 0.93 (Range: 0.75 – 1.47). The median value of relative ratio in *mm* vs *fm* was 0.90 (Range: 0.41 – 1.81), *mm* vs *ff was* 1.07 (Range: 0.55 – 2.93), *mf* vs *fm* was 0.99 (Range: 0.32 – 1.25), *mf* vs *ff* was 1.17 (Range: 0.43 – 3.21) and *ff* vs *fm* was 0.75 (Range: 0.35 – 1.06) (Figure 9).

**Figure 8.**
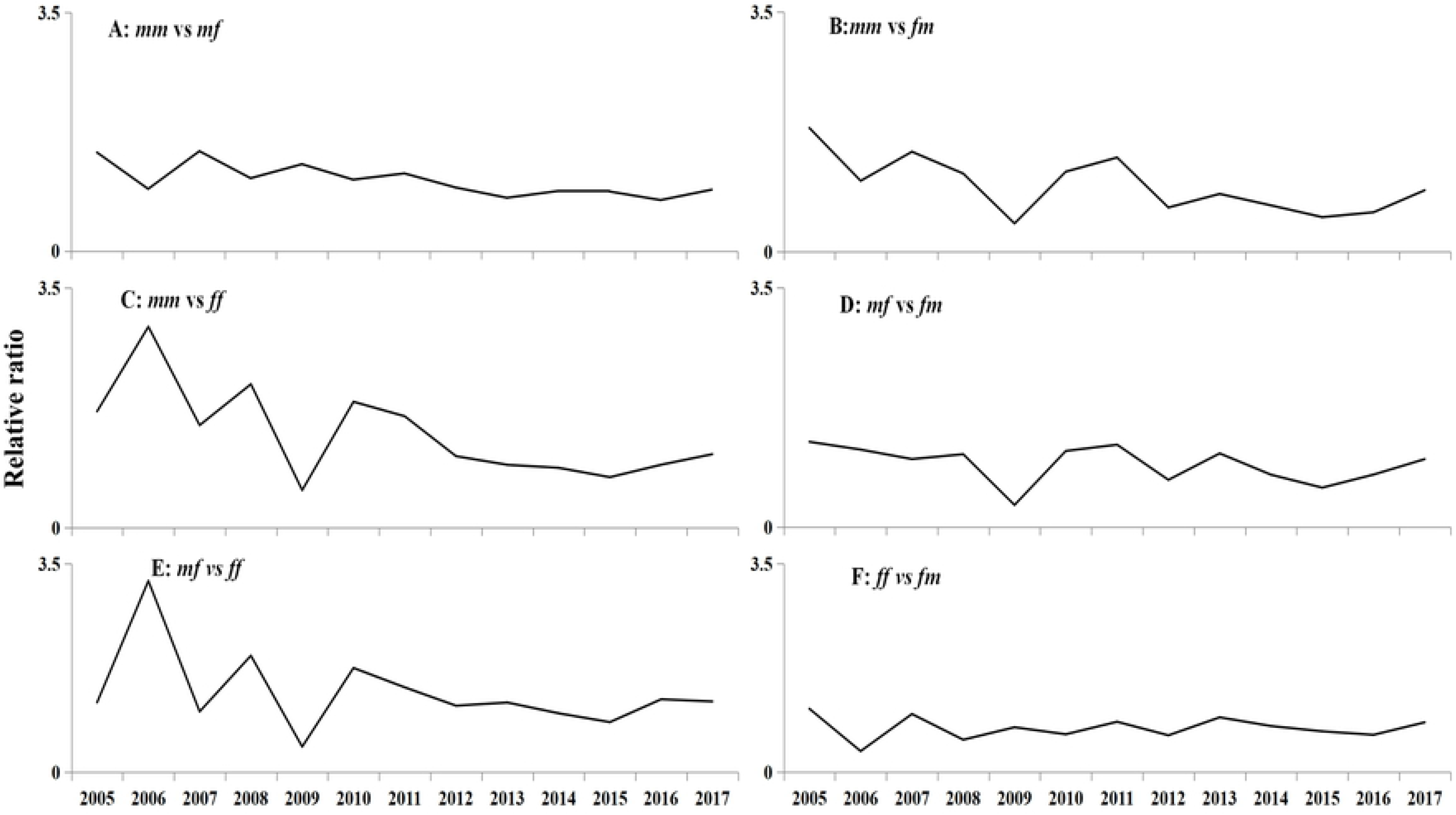
The five kinds of relative ratio from 2005 to 2017 in Hubei (A: *mm* vs *mf*, among male versus from male to female; B: *mm* vs *fm*, among male versus from female to male; C: *mm* vs *ff*, among male versus among female; D: *mf* vs *fm*, from male to female versus from female to male; E: *mf* vs *ff*, from male to female versus among female; F: *ff* vs *fm*, among female versus from female to male).

**Figure 9.**
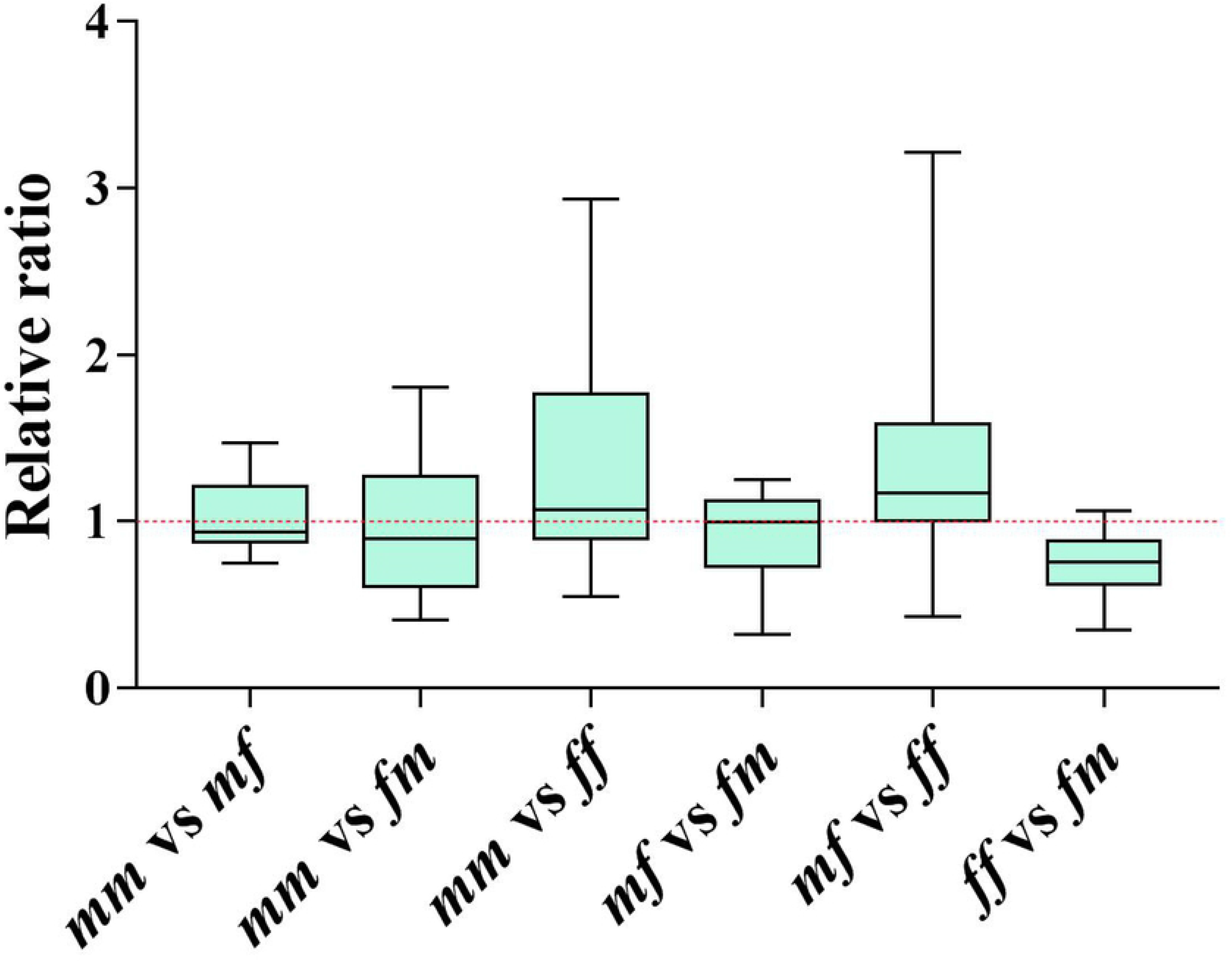
Box-plot of relative ratio from 2005 to 2017 in Hubei (*mm:* among male; *ff:* among female; *fm*: from female to male; *mf*: from male to female).

## Discussion

In this study, we were the first ones to make the transmission of shigellosis between different genders clear. We applied SEIAR model to explore the differences of the water-borne infectious disease in males and females for the first time. It has guiding significance for controlling the prevalence of shigellosis.

### Validity of the model

According to *R^2^* of linear regression, the model of SEIAR has a high good-of-fitness with the reported data in different genders. It is consistent with the results of a research [6], suggesting the model is suitable for this study.

### Epidemiological characteristics

In recent years, although the incidence of shigellosis has a decreasing trend in China [16, 21, 22], it is still relatively high level in Hubei province from 2005 to 2017. The difference incidence of shigellosis cases in male and female is observed by the descriptive epidemiology [23, 24]. However, all of them do not clarify the reasons for the difference. A study indicates that there were more male than female cases (the ratio of male to female is 1.3:1), which is consistent with our results in descriptive epidemiology [25].

According to a new review[1], the transmission pattern of shigellosis has shifted from water/food-to-person to from person-to-person, with high risk groups being particularly men who have sex with other men in developed country. Does this mean that the transmissibility of shigellosis among males is stronger than among females? We developed SEIAR model to verify this hypothesis. However, we obtained the number of cases in five hypotheses using “knock-out” simulation. When *β_fm_* =0, the number of cases dropped most in different genders, which means that the female-to-male had a large contribution during the transmission. It is important to isolate and treat female cases, and to strengthen the personal health.

### Transmissibility of Shigellosis in different genders

Compared with HIV which has different transmissibility in different genders, shigellosis is not particularly highly contagious in different genders [26]. Our results showed that the mean values of the transmission parameters among males and females, from male to female, and from female to male are different, and they have the following order: *β_fm_* > *β_m_* > *β_mf_* > *β_f_*. The median values of *SAR* have the following order: *SAR_fm_* > *SAR_mf_* > *SAR_mm_* > *SAR_ff_*. The median values of relative ratio of *SAR* have the following order: *mf* vs *ff* > *mm* vs *ff* > *mf* vs *fm* > *mm* vs *mf* > *mm* vs *fm* > *ff* vs *mf*. All the results have a common feature, that the transmission is mainly female-to-male. These findings showed that male individuals are more transmissible than female individuals. Therefore, the different transmissibility between males and females is the reason for the difference in distribution between genders.

There are a large number of studies focused on the distribution of incidence in different age groups [1, 7, 16, 22]. And the high-risk group is under 5 years old and over 60 years old. Combining with our results and the actual situation of China, whether it can be considered related to the tradition of the elderly bringing children at home. Elderly people, especially grandmothers, have more daily contact with their children, which lead to such high transmission rates. Meanwhile, the transmissibility of shigellosis in different age groups further needs to be studied.

### Limitation

According to a recent study, although it is mainly transmitted from person to person [1], the shigellosis is still a water/food-borne disease. For this reason, there has been an impact on our result given that we simplified the SIEARW model and ignored environmental factors (water and food). At the same time, the parameters of SIEAR model come from relevant references and Hubei Statistical Yearbook, not from collection, this has an impact on the accuracy of our model.

### Conclusions

In Hubei Province, The incidence of shigellosis in males is higher than that in female, causing the disease to be a burden. The transmissibility of shigellosis is different in male and female individuals. Males seem to be more transmissible than females and the transmission is mainly female-to-male.

## Data Availability Statement

All relevant data are within the paper and its Supporting Information files.

## Acknowledgments

We thank the staff members in the hospitals, local health departments, and local CDCs for their valuable assistance in coordinating data collection.

## Supplementary Information Legends

**S1 Table. Goodness of fit of curve fitting in different sub-seasons in each year from 2005 to 2017 in Hubei Province, China**

